# Timing and causes of the distribution pattern of *Oncomelania hupensis* estimated by molecular and geologic data

**DOI:** 10.1101/450031

**Authors:** Wen-bin Ji, Shu-xin Xu, Jun Bai, Ying-yi Cui, Xian-min Zhou, Jie-xin Zou

## Abstract

As the only intermediate host of *Schistosoma japonicum*, *Oncomelania hupensis* plays an irreplaceable role in the prevalence of schistosomiasis japonica. Several living subspecies of *Oncomelania hupensis* are found in Asia, especially in China, Japan,the Philippines, and Sulawesi of Indonesia. The existing geographical distribution pattern of *O. hupensis* has been influenced by geological events. This study used existing mitochondrial gene data for *O. hupensis* in the GenBank database and the molecular clock method to estimate the divergence time of each subspecies of *O. hupensis*. For the first time, the timing and causes of the distribution pattern of the different *O. hupensis* subspecies were studied by combining molecular data with data on geologic events. The results showed that the uplift and isolation of the Qinghai-Tibet Plateau caused *Oncomelania hupensis robertsoni* to differentiate 4.76 Ma(Million anniversary), while *Oncomelania hupensis guangxiensis* was affected by the third Himalayan orogenic movement, differentiating 1.10 Ma. *Oncomelania hupensis nosophora* was influenced by the formation of the Yonaguni Strait and diverged 1.43 Ma. Influenced by ice ages and interglacial periods, *Oncomelania hupensis tangi* and *Oncomelania hupensis formosana* diverged 0.57 Ma. The link of the ancient Yangtze River promoted the spread of *O. hupensis* to the middle and lower reaches of the Yangtze River, and the developed water network facilitated gene exchange among *Oncomelania hupensis hupensis* in the area. Eventually, 0.62 Ma, *O. h. hupensis* differentiated.

**Author summary:** Phylogenetic analysis of Pomatiopsidae species showed that *Oncomelania* was isolated from other genera and clustered independently in phylogenetic trees. Further analysis of the species *Oncomelania hupensis* and its subspecies was performed. The snail species *O. hupensis* has multiple subspecies that exhibit certain differences. These subspecies are distributed across Asia, from China’s Yunnan Province in the west to Japan in the east and south to the Philippines and Indonesia. In addition, the subspecies are widespread in the middle and lower reaches of the Yangtze River in China, and the distribution ranges of the different subspecies do not overlap. The formation of this distribution pattern of each subspecies of *O. hupensis* has a profound impact on the prevalence of *Schistosoma japonicum*. Therefore, the authors analyzed molecular data and geological historical events to investigate the timing and causes of the distribution pattern of each subspecies of *O. hupensis*.

## Introduction

*Oncomelania hupensis* (Gredler, 1881) are dioecious, amphibious freshwater snails belonging to the phylum Mollusca, the class Gastropoda, the order Mesogastropoda, and the family Pomatiopsidae (Stimpson, 1865). *Oncomelania* (Gredler, 1881), as the only intermediate host of *Schistosoma japonicum* (Katsurada, 1904), plays an important role in the spread of *S. japonicum* [1, 2] and is now located in China, Japan, the Philippines and Indonesia [3].

As the only intermediate host of *Schistosoma japonicum*, the control and killing of *Oncomelania hupensis* will be the key link to block the transmission of schistosomiasis japonica. At present, the control and prevention of schistosomiasis japonica have made great progress [2, 4, 5], but researchers are still searching for more effective methods and tools [6, 7] to eliminate the prevalence of schistosomiasis japonica in the end [8]. The distribution of *Oncomelania hupensis* is overlapping with the endemic area of schistosomiasis japonica, and the transmission of *Oncomelania hupensis* will affect the prevalence of schistosomiasis japonica to some extent [9], so this paper thinks that the study of the distribution of *Oncomelania hupensis* and the cause and time of distribution pattern formation will be of significance to control the prevalence of schistosomiasis japonica.

Based on morphological data, G.M. Davis et al. [10] performed the first systematic classification of *Oncomelania* species. *Oncomelania* includes two species, *Oncomelania.minima* (Bartsch, 1936) and *O. hupensis. O. hupensis* comprises a total of six subspecies, including *Oncomelania hupensis chiui* (Habe & Miyazaki, 1962), *Oncomelania hupensis formosana* (Pilsbry & Hirase, 1905), *Oncomelania hupensis lindoensis* (Davis & Carney, 1973), *Oncomelania hupensis nosophora* (Robson, 1915), *Oncomelania hupensis quadrasi* (Möellendorff, 1895) and *Oncomelania hupensis hupensis*. Based on morphological classification, Yueying Liu et al. [11] suggested that the *O. hupensis* distributed on the Chinese mainland can be divided into five subspecies. Through molecular genetic analysis combined with morphological analysis, Yibiao Zhou et al. [12] proposed the classification of four subspecies of *O. hupensis*: *Oncomelania hupensis robertsoni* (Bartsch, 1936), *Oncomelania hupensis tangi* (Bartsch, 1936)*, Oncomelania hupensis hupensis*, and *Oncomelania hupensis guangxiensis* (Liu et al., 1981). In the process of exploring novel microsatellite markers of *O. hupensis*, Xiaonong Zhou et al. [13] found that *O. hupensis* on mainland China exist in four different ecological environments: swamps and lakes in the Yangtze River basin, mountains and hills in the Sichuan and Yunnan provinces, coastal areas in Fujian Province, and karst landforms in the Guangxi autonomous region, which is consistent with the classification of Zhou Yibiao et al.

The existing *O. hupensis* subspecies are distributed in Asia with a geographically isolated distribution pattern. There is no overlap of distribution [14], and there are different hypotheses that suggest that this distribution pattern is due to the spread and differentiation of *O. hupensis* and the influence of geological plate movement.

Hypothesis one: Based on morphological data, G.M. Davis et al. [10, 15] constructed the Pomatiopsidae phylogenetic atlas and an evolutionary tree, examined the distribution of the snails in river basins in Asia, and proposed that the snail ancestors lived on the Indian continental plate during the Mesozoic period. Davis further proposed that during the Himalaya orogeny, the snails entered new Yangtze River waters in the north of Mengmi and Yunnan and spread over the Pacific coast of China and to Japan; Taiwan Province; the Philippines; and Sulawesi in Indonesia. In 1995, Xiaonong Zhou et al. [16] proposed that the snails entered China’s Yunnan Province earlier from Himalaya, then moved into the Sichuan plain and spread to the eastern coast of mainland China.

Hypothesis two: S.W. Attwood [17] thinks that proto-Pomatiopsidae came from northwest Australia (during the Jurassic period) on small continental fragments of Gondwanan origin. The continental fragments were involved in the formation of present-day Borneo and eastern Indonesia. During the Oligocene, the proto-Pomatiopsidae began to diverge from Borneo and eastern Indonesia, spreading north to Japan before the opening of the Sea of Japan and colonizing China and the mountains of western China.

The hypotheses above are based on speculated tectonic plate movement, and there are some limitations. This study used GenBank data and bioinformatics methods, fossil ages, and the molecular clock method [18] to estimate the divergence times of existing *O. hupensis* subspecies. The divergence of the existing *O. hupensis* subspecies is discussed in the context of historical geological events. Thus, molecular data and information on geological events were used to verify the above hypotheses and to elucidate the migration and subspeciation of these snails.

## Data and methods

### Data

In the GenBank database, there were 26 mitochondrial genome sequences for *O. hupensis* in Asia (S1 Table), including those for the subspecies *O. h. nosophora*, *O. h. quadrasi*, *O. h. hupensis* and *O. h. robertsoni*. In addition, *O. h. chiui* had 11 COI gene sequences, and *O. h. formosana* had 3 COI gene sequences and 1 16S gene sequence, although these subspecies did not have mitochondrial gene sequences. No molecular data for *O. h. lindoensis* had been uploaded.

### Methods

#### Phylogenetic analysis based on 13 protein-coding genes (PCGs)

*Tricula hortensis* (EU440735) was chosen as the outgroup. Mafft [19] software was used to compare the sequences, and the Gblocks Server (http://molevol.cmima.csic.es/castresana/Gblocks_server.html) was used to select the conserved sequence area. The best-fitting evolutionary model in maximum likelihood (ML) analyses of the 13 PCGs was GTR+G+I, and it was selected using ModelGenerator [20]. This model was used in Mega 6.0 [21], and 1,000 bootstrap replicates were performed. For the Bayesian inference (BI) analyses, the best-fitting evolutionary model (GTR+G+I) was determined using MrModeltest 2.3 [22].

The BI analyses were performed with MrBayes 3.2.6 [23], which used the Markov chain Monte Carlo (MCMC) method with four chains to run 2 million generations. Sampling was performed every 1,000 generations, and the initial 500,000 steps were discarded as burn-in. The consensus BI trees were visualized using FigTree 1.4.0 software.

#### Divergence time estimation based on molecular clock

Upon searching for fossil ages in Gastropoda, we obtained the fossil age of the family Assimineidae (23 Ma-16 Ma) [24, 25]. In the GenBank database, there are sequences for COI genes and 16S mitochondrial genes for four species in Assimineidae: *Angustassiminea satumana* (AB611803/AB611802), *Assiminea hiradoensis* (AB611807/AB611806), *Paludinellassiminea japonica* (AB611811/AB611810), and *Pseudomphala miyazakii* (AB611815/ AB61181). The fossil ages and gene sequences were used to estimate the subsequent fossil ages. *O. minim* also belongs to the genus *Oncomelania*, and *O. minima* had COI and 16S single gene data in the GenBank database (AB611791/AB611790, AB611795/AB611794, and DQ212795/DQ212858). *O. h. formosana* also had COI and 16S gene sequences (DQ112283/DQ212861), so *O. minima* and *O. h. formosana* were included in the estimation of divergence time. *O. h. chiui* and *O. h. lindoensis* were not included in the divergence time estimate. To further improve the reliability of the divergence time estimation, GenBank COI and 16S gene sequences (S2 Table) from the same individual source in the 14 genera and 31 species of Pomatiopsidae were also included.

We used the strict clock in BEAST v1.8.2 [26] software to estimate the divergence time of each subspecies based on the COI and 16S sequences and selected a normal prior distribution. The best-fitting evolutionary model in ModelGenerator [20] software was GTR+G+I, and the Yule process was used for the tree prior with a random starting tree. In total, 20,000,000 generations were run, and the parameters were logged every 20,000 generations, with the first 10% discarded as burn-in. TreeAnnotator v1.8.2 software was used to generate the tree, and the divergence time was visualized using FigTree 1.4.0 software.

## Results and discussion

### Phylogenetic relationships and classification

The occurrence tree showed that 26 of the *O. hupensis* sequences were grouped into 6 branches according to subspecies(Fig 1). The 6 subspecies were *O. h. hupensis*, *O. h. tangi*, *O. h. guangxiensis*, *O. h. nosophora*, *O. h. quadrasi* and *O. h. robertsoni*, which conform to the subspecies classification system proposed by G.M.Davis [10,27] and Yibiao Zhou [12].

**Fig. 1:**
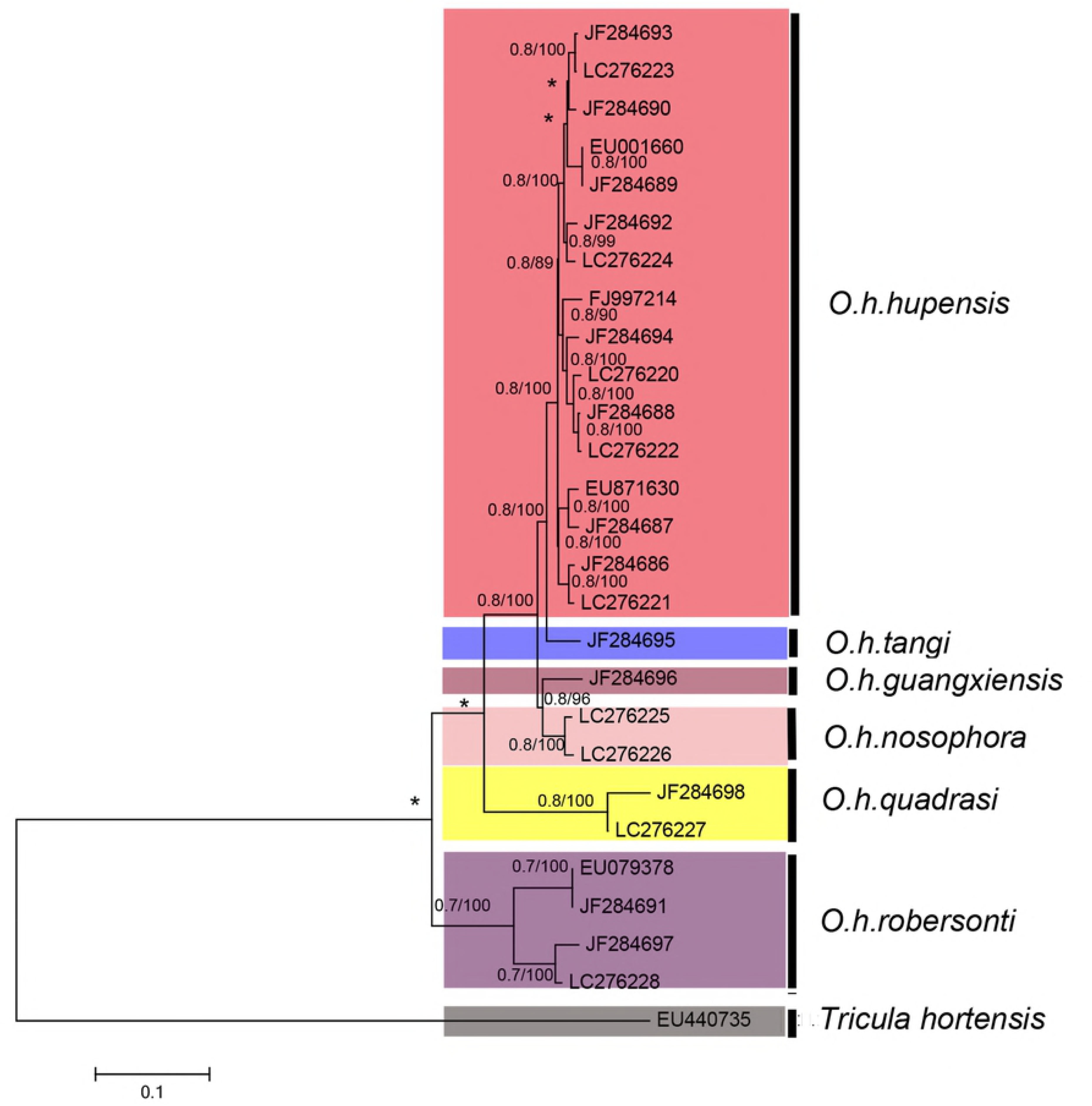
Maximum likelihood (ML) phylogenetic tree of *Oncomelania hupensis* and related brachyurans based on 13 PCG sequences from the mitochondrial genome. *Tricula hortensis* serves as the outgroup. The numbers at the internodes are ML bootstrap probabilities and Bayesian inference (BI) posterior probabilities. The differences between the ML and BI trees are indicated by‘*’. The scale bars represent the genetic distance.

### Divergence time estimation

The existing research results show that the subfamily Triculinae of Pomatiopsidae is mainly distributed in Southeast Asia, and the differentiation of the species is closely related to the temporal variation of the geologic events in the distribution area [28], which are closely related to the results of this paper. Although *Oncomelania* is now classified as part of Pomatiopsidae, the molecular clock (Fig 2) estimates show that *Oncomelania* species diverged at 12.41Ma,earlier than the rest of the included species in the family, suggesting the independence of *Oncomelania*. This paper will focus on the differences in the divergence times of *Oncomelania* species and on the divergence events.

**Fig. 2:**
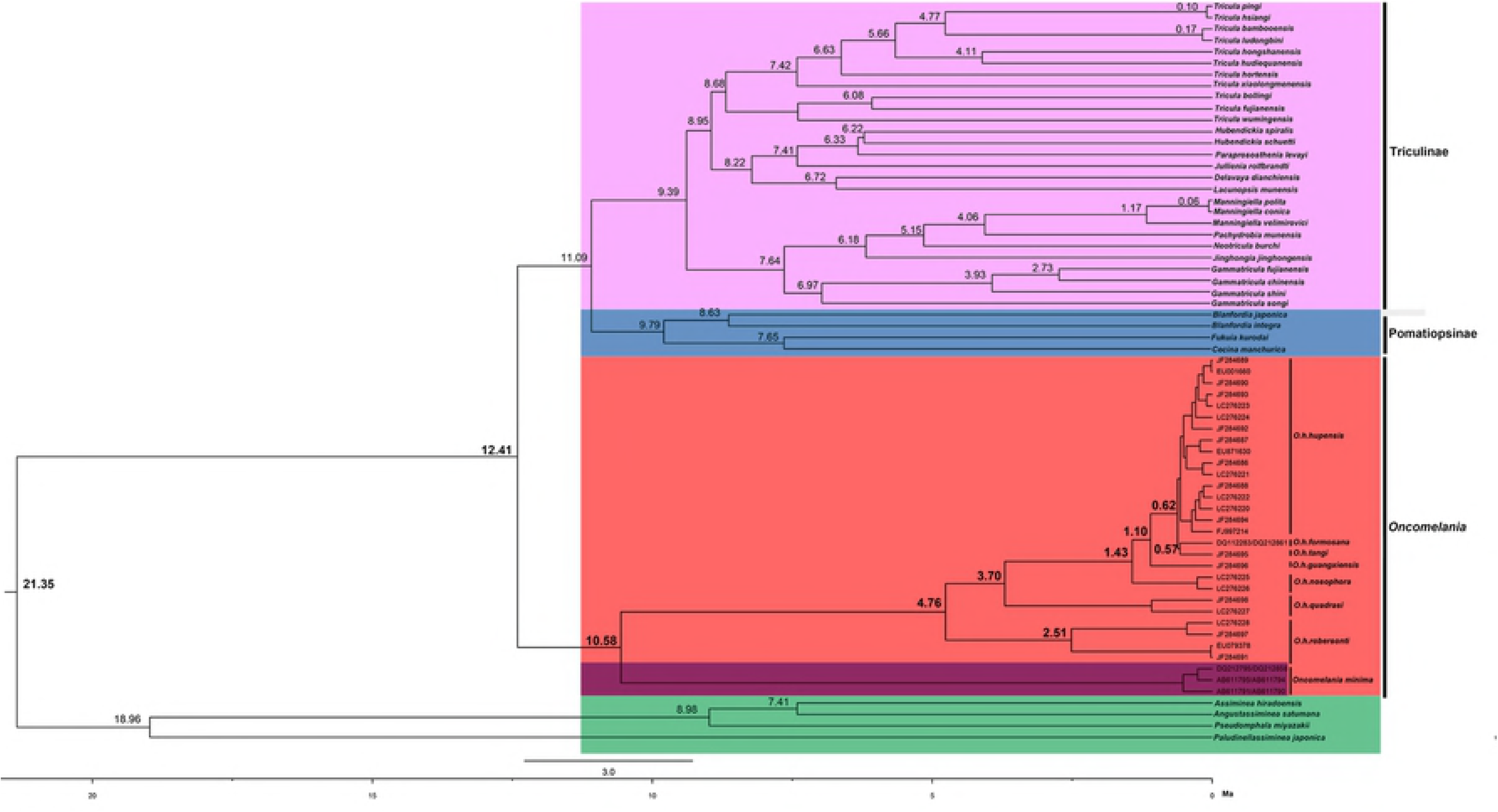
Divergence time of Pomatiopsidae based on the COI and 16S sequences of the mitochondrial genome. The scale bars represent millions of years from the present.

Combining the molecular clock calculation results based on the COI and 16S gene sequences and corresponding periods of geological events, this article on the timing and cause of *Oncomelania* species (subspecies) migration and differentiation makes the following speculation (Fig 3):

**Fig. 3:**
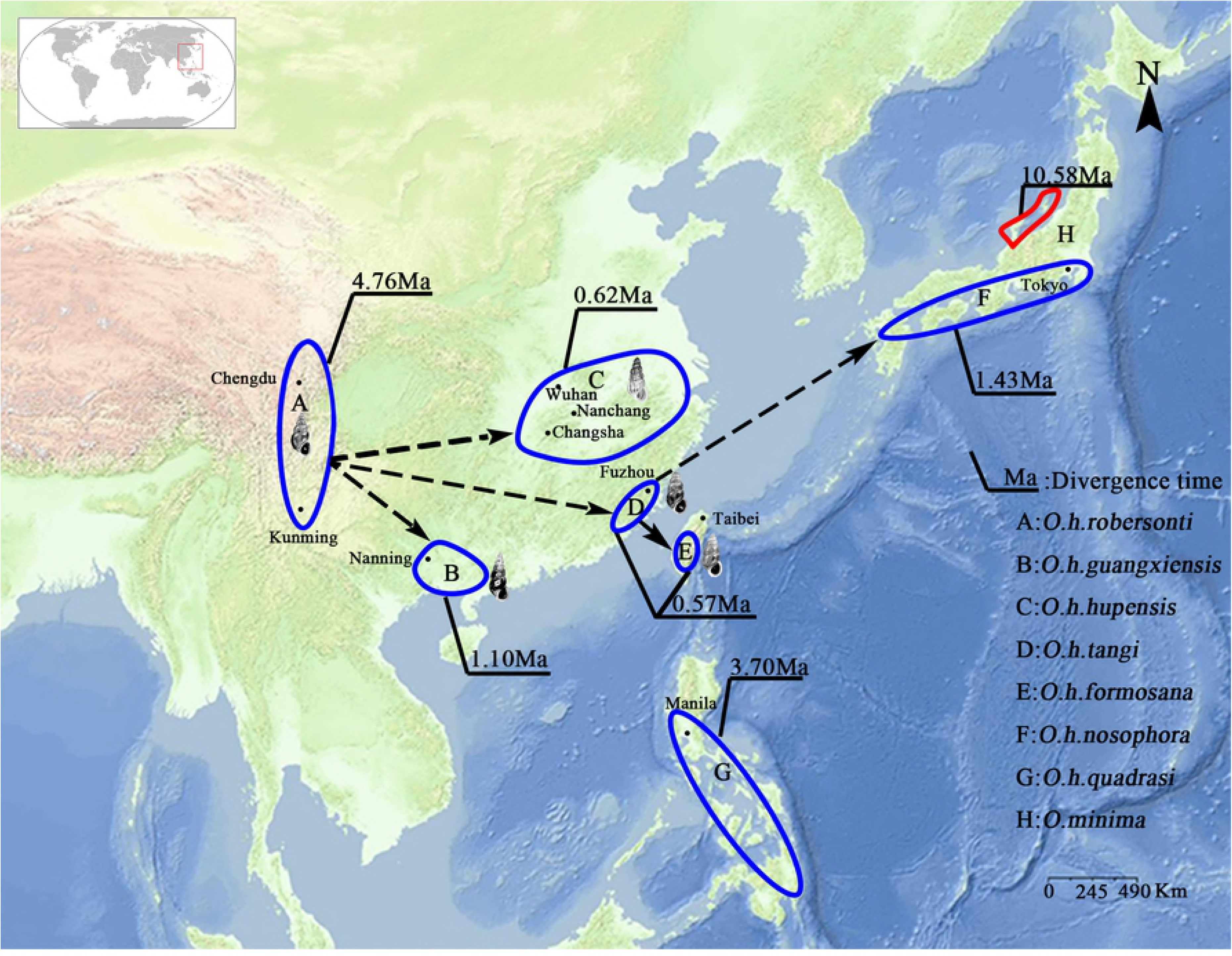
Distribution and differentiation of *Oncomelania* in Asia. The regional map of Asia comes from https://commons.wikimedia.org/wiki/Atlas_of_the_world and http://landsatlook.usgs.gov/; the map edited with Adobe Photoshop CS6. Ma (Million anniversary).

The ancestors of *O. hupensis* were distributed on the Asian continent. During the period from 23 Ma to 15 Ma [29], the Sea of Japan opened, and the Japanese islands gradually separated from the Asian continent, resulting in geographical isolation and a new ecological environment. Thus, 10.58 Ma (95% confidence interval = 7.34-14.07 Ma), *O. minima* diverged. This may be a major cause of the current distribution of *O. minima* in Honshu, Japan, while the snails on the Asian continent are differentiated into different subspecies of *O. hupensis*.

On the Asian continent, approximately 60-50 Ma [30], the Indian Ocean Plate and the Eurasian Plate began to collide and cause the Qinghai-Tibet Plateau to uplift. The uplift of the Qinghai-Tibet Plateau was not a rapid process but rather proceeded through different stages. After the plate collision, the strong uplift began 3.4 Ma [31]. The results show that the divergence time of *O. h. robertsoni* was 4.76 Ma (95% confidence interval = 2.44-4.98 Ma), which is close to the stage of strong uplift. With the elevation of the Qinghai-Tibet Plateau, the change in climate [32], the change in ecological environment, and the development of geographical isolation, the snail species gradually differentiated in this location, producing *O. h. robertsoni*. In addition, we found that the divergence time of *S. japonicum* was approximately 3.8 Ma [33, 34], close to the divergence time of *O. h. robertsoni*, which may represent the beginning of the parasitic relationship between *S. japonicum* and *O. hupensis* [35]. When *S. japonicum* began to live in the snail body and whether there was a symbiotic relationship between the species remains to be elucidated.

The Quaternary glaciation (2.58 Ma) has included four ice ages (1.50-1.10 Ma, 0.90-0.40 Ma, 0.20-0.11 Ma, and 0.01-0.00 Ma) [36, 37] and interglacial periods. The results of this study show that the divergence time of *O. h. formosana* in Taiwan Province and *O. h. tangi* in Fujian Province was 0.62 Ma (95% confidence interval = 0.40-0.85 Ma), which coincides with the separation of the island of Taiwan from the continent due to the transition of an ice age into an interglacial period. There is reason to believe that the formation of *O. h. formosana* and *O. h. tangi* was affected by such a transition. The island of Taiwan and the mainland were connected during the ice age, and the *O. hupensis* in the two places were able to carry out gene exchange. During the interglacial periods, the island of Taiwan was again separated from the mainland, reestablishing geographical isolation, and *O. h. formosana* and *O. h. tangi* eventually diverged.

The formation of the Yonaguni Strait 1.55 Ma led to the separation of the island of Taiwan from the Japanese islands [38]. The results suggest that the divergence time of *O. h. nosophora* in Japan was 1.43 Ma (95% confidence interval = 0.90-2.00 Ma), which coincides with the separation of the island of Taiwan from the islands of Japan. It is reasonable to assume that the geographical isolation caused by the strait facilitated the formation of *O. h. nosophora.*

In this study, the divergence time of *O. h. guangxiensis* was found to be 1.10 Ma (95% confidence interval = 0.70-1.55 Ma). Guangxi is located on the southeast edge of the Qinghai-Tibet Plateau and was affected by the third Himalayan orogenic movement (starting 2.4 Ma) [39, 40]; geological activity and further geologic changes produced geographic isolation and eventually resulted in the differentiation of *O. h. guangxiensis*.

The uplift of the Qinghai-Tibet Plateau caused the ancient Yangtze River to penetrate from west to east during the period from 2.6 to 2.0 Ma [41]. The developed water network of the ancient Yangtze River basin strongly aided the spread of the snails to the middle and lower reaches of the Yangtze River, which had their own network and provided a broad living space for the multiplication and differentiation of *O. hupensis*, thus forming the named subspecies *O. h. hupensis* as the largest group. The changes in the Yangtze River coincide with the divergence time of this subspecies, 0.62 Ma (95% confidence interval = 0.40-0.85 Ma).

The divergence time of *O. h. quadrasi* was 3.70 Ma (95% confidence interval = 2.44-4.98 Ma), and there are no corresponding geological events to explain the cause of the differentiation. However, according to the two factors of *O. h. robertsoni* and the carrying of *S. japonicum*, it is reasonable to infer that *O. h. quadrasi* originated from *O. h. robertsoni*.

## Conclusion

Through analysis of the molecular clock results and of molecular data and geologic events, the timing and causes of the distribution pattern of the different *O. hupensis* subspecies were studied for the first time. We speculate that the isolation of the Japanese islands from the mainland caused *O. minima* to differentiate 10.58 Ma and that the isolation caused by the uplift of the Qinghai-Tibet Plateau 4.76 Ma caused *O. h. robertsoni* to differentiate. Upon the west-to-east formation of the river, geological changes and changes in the river influenced the beginning of the differentiation of the various existing subspecies of *O. hupensis*. Among them, *O. h. guangxiensis* was affected by the third Himalayan orogenic movement and differentiated 1.10 Ma. Influenced by the formation of the Yonaguni Strait, *O. h. nosophora* differentiated 1.43 Ma, while 0.57 Ma, *O. h. tangi* and *O. h. formosana* diverged under the influence of four ice ages and interglacial periods. The link of the ancient Yangtze River promoted the spread of *O. hupensis* to the middle and lower reaches of the Yangtze River, and the developed water network facilitated the gene exchange of *O. hupensis* in the area; the most numerous and most widely distributed subspecies, *O. h. hupensis*, diverged 0.62 Ma.

## Acknowledgments

The authors thank the guidance provided by Xiaonong Zhou, Director of the National institute of parasitic diseases Chinese Center For Disease Control And Prevention (NIPD) for this article.

## Supporting information

**S1 Table. Mitochondrial genome data for Oncomelania hupensis in GenBank (as of August 7, 2018)**. *indicates that the sequence upload did not add the sample source to GenBank.

**S2 Table. Partial COI and 16S genes of 14 genera and 31 species in Pomatiopsidae from GenBank (as of August 7, 2018)**. *indicates that the sequences are from the same mitochondrial genome.

